# Whole genome sequencing reveals high complexity of copy number variation at insecticide resistance loci in malaria mosquitoes

**DOI:** 10.1101/399568

**Authors:** Eric R. Lucas, Alistair Miles, Nicholas J. Harding, Chris S. Clarkson, Mara K. N. Lawniczak, Dominic P. Kwiatkowski, David Weetman, Martin J. Donnelly, The Anopheles gambiae 1000 Genomes Consortium

**Affiliations:** Liverpool School of Tropical Medicine, Pembroke Place, Liverpool L3 5QA, UK.; Wellcome Sanger Institute, Hinxton, Cambridge CB10 1SA, UK.; Big Data Institute, University of Oxford, Li Ka Shing Centre for Health Information and Discovery, Old Road Campus, Oxford OX3 7LF, UK.

**Keywords:** CNV, gene duplication, metabolic insecticide resistance, *Anopheles*, whole genome sequencing, Ag1000G

## Abstract

**Background:** Polymorphisms in the copy number of a genetic region can influence gene expression, coding sequence and zygosity, making them powerful actors in the evolutionary process. Copy number variants (CNVs) are however understudied, being more difficult to detect than single nucleotide polymorphisms. We take advantage of the intense selective pressures on the major malaria vector *Anopheles gambiae*, caused by the widespread use of insecticides for malaria control, to investigate the role of CNVs in the evolution of insecticide resistance.

**Results:** Using the whole-genome sequencing data from 1142 samples in the *An. gambiae* 1000 genomes project, we identified 1557 independent increases in copy number, encompassing a total of 267 genes, which were enriched for gene families linked to metabolic insecticide resistance. The five major candidate genes for metabolic resistance were all found in at least one CNV, and were often the target of multiple independent CNVs, reaching as many as 16 CNVs in *Cyp9k1*. These CNVs have furthermore been spreading due to positive selection, indicated by high local CNV frequencies and extended haplotype homozygosity.

**Conclusions:** Our results demonstrate the importance of CNVs in the response to selection, with CNVs being closely associated with genes involved in the evolution of resistance to insecticides, highlighting the urgent need to identify their relative contributions to resistance and to track their spread as the application of insecticide in malaria endemic countries intensifies. Our detailed descriptions of CNVs found across the species range provides the tools to do so.

## 1 Introduction

Copy number variants (CNVs) are a form of genetic variation that occur when a genomic sequence is deleted or duplicated, potentially affecting both the structure and expression levels of coding sequences and playing a crucial role in evolution and adaptation [1-3]. Part of the importance of CNVs lies in the wide range of effects that they can have on the transcriptome. Increases in copy number (amplifications) encompassing the entire sequence of a gene can lead to elevated expression levels if new gene copies are associated with cis-regulatory sequences required for transcription [4]. Alternatively, duplication or deletion of only part of a gene’s sequence can lead to structural variation. For example, in humans, a CNV spanning parts of two glycophorin genes creates a novel hybrid glycophorin associated with resistance to malaria [5]. CNVs can also allow alternative variants of a gene to appear in tandem on the same chromosome through heterogeneous gene duplication, creating constitutive heterozygotes. This can be seen in the mosquitoes *Anopheles gambiae* and *Culex pipiens,* where mutations in Acetylecholinesterase-1 (*Ace-1*) cause resistance to carbamate and organophosphate insecticides, but carry a fitness cost in the absence of insecticide. This cost is mitigated in heterozygotes, leading to the spread of heterogeneous *Ace-1* duplications in which the mutant and wild-type alleles co-occur [6,7].

While the importance of CNVs is widely recognised, they typically receive less attention than single nucleotide polymorphisms (SNPs) in investigations of genetic variation, perhaps because they are more difficult to identify. Population-level genome-wide analyses of CNVs are thus rare and the extent of their impact on evolution is poorly understood (but see [8] for a worldwide study in humans).

The malaria mosquito *An. gambiae* and its close sister species *An. coluzzii* provide excellent models to study the evolution of CNVs at a population level for three reasons. First, these species are the major vectors of malaria in Sub-Saharan Africa (SSA) and are highly anthropophagic [9]. As a consequence, they are heavily targeted by insecticides used in malaria control programmes, creating intense selective pressures that drive rapid contemporary evolution. CNVs thus have an opportunity to contribute to the selective response to these pressures, providing a context in which their importance can be assessed. Second, CNVs can play a key role in the evolution of insecticide resistance due to their ability to affect gene expression and allow co-expression of wild-type and mutant alleles [10,11]. Despite nearly two decades of genetic research into insecticide resistance, known resistance-associated SNPs are still unable to explain much of the variance in insecticide resistance [12]. CNVs potentially represent a crucial source of missing variation that can fill this gap. Third, the *Anopheles gambiae* 1000 Genomes project (Ag1000G) has produced whole-genome sequencing data from 1142 individual *An. gambiae* and *An. coluzzii* from multiple locations in SSA, providing a unique opportunity to conduct genome-wide searches for CNVs from across the species’ distributions [13].

The two major mechanisms of insecticide resistance are target site resistance and metabolic resistance [14]. CNVs have been found to affect all three major insecticide target site genes in insects (*Ace-1*: [6,15]; the *para* voltage-gated sodium channel (Vgsc): [16,17]; gamma-aminobutyric acid (GABA) receptor: [18]), usually combining resistant and wild-type alleles to provide resistance while mitigating its cost. However, duplications in *Ace-1* are the only CNVs so far shown to play a role in *Anopheles* insecticide resistance, being associated with a resistance-conferring mutation and either increasing the resistance which it confers [19] or diminishing the fitness cost of the mutation [6].

Metabolic genes whose expression levels are associated with insecticide resistance have been reported in a wide range of species [20], but the causative genetic variants often remain unidentified. Focused studies have identified cases where CNVs play a critical role in metabolic insecticide resistance in a range of species. In *Drosophila*, duplication of the detoxification gene *Cyp6g1* has been implicated in resistance to DDT [21], while in *Cx. quinquefasciatus* resistance to permethrin is associated with increased expression of *Cyp9m10*, due in part to a duplication [22] Similarly, amplification of esterase genes leading to elevated expression provides increased resistance to organophosphates in the mosquitoes *Cx. pipiens* [23] and *Aedes albopictus* [24], and to several insecticides in the peach-potato aphid *Myzus persicae* [25,26]. In brown planthoppers, neofunctionalisation of a duplicated copy of *Cyp6er1* has even created a novel gene variant providing resistance to the neonicotinoid imidacloprid [27]. In *An. gambiae* and *An. coluzzii,* the most import metabolic genes that have been identified as major insecticide resistance candidates to 94 date are *Gste2* [28], *Cyp6p3* [29,30], *Cyp6m2* [29,31,32], *Cyp6z1* [33] and *Cyp9k1* [34,35]. If CNVs play an important role in the evolution of insecticide resistance in *An. gambiae* and *An. coluzzii*, we would expect to find them among genes such as these.

Here, we use the whole-genome sequencing data resources of the Ag1000G project to perform an agnostic genome-wide scan of CNVs in *An. gambiae* and *An. coluzzii*. We find that CNVs are enriched for genes and gene families implicated in insecticide resistance, with many independent CNV events encompassing the five key metabolic resistance-associated genes (*Gste2*, *Cyp6p3*, *Cyp6m2*, *Cyp6z1* and *Cyp9k1*). Further investigation into these genes reveals that several CNVs exist at high local frequencies and show clear evidence of strong positive selection.

## 2 Results

We found a total of 1557 independent CNVs in the Ag1000G phase 2 dataset using sequencing coverage data (Supplementary Data S1); 250 of these CNVs contained at least one gene, significantly more than expected by chance (*P* < 0.0001, null distribution determined by simulation). Out of 10,939 genes included in the analysis, 267 (2.4%) were found in at least one CNV (Supplementary Data S2). As expected, these included the well-documented duplication in the insecticide target site *Ace-1*. No CNVs were found in the other common insecticide target sites Vgsc or GABA.

### 2.1 CNVs are enriched for potential metabolic resistance genes

Of the 267 genes found in CNVs (Supplementary Data S2), 28 were candidate metabolic resistance genes (defined as a cytochrome P450, glutathione-S-transferase or carboxylesterase, and referred to as “metabolic detox genes” from here on). Because many related genes occur in clusters, and are therefore not independently included in CNV events, we counted the number of CNVs that included at least one metabolic detox gene. Of the 250 CNVs that contained any genes, 27 contained at least one metabolic detox gene, significantly more than expected by chance (*P* < 0.0001, null distribution determined by simulation). While there was some variation between populations in the number of metabolic detox genes found in CNVs (Table 1), this was not significant (Fisher’s exact test: *P* = 121 0.08).

**Table 1:** Number of CNVs detected either containing or not containing at least one cytochrome P450, glutathione-S-transferase or carboxylesterase (“detox genes”). AO = Angola, BF = Burkina Faso, CI = Côte d’Ivoire, CM = Cameroon, FR = French Mayotte, GA = Gabon, GH = Ghana, GM = The Gambia, GN = Guinea, GQ = Equatorial Guinea, GW = Guinea-Bissau, KE = Kenya, UG = Uganda. col = *An. coluzzii*, gam = *An. gambiae*. Numbers in brackets after the population name indicate the total number of samples from that population after removal of high-variance samples.

|  | AOcol<br>(68) | BFcol<br>(75) | BFgam<br>(91) | CIcol<br>(71) | CMgam<br>(297) | FRgam<br>(23) | GAgam<br>(68) | GHcol<br>(55) |
| --- | --- | --- | --- | --- | --- | --- | --- | --- |
| CNVs without detox genes | 377 | 330 | 342 | 365 | 335 | 215 | 447 | 304 |
| CNVs with detox genes | 4 | 5 | 5 | 7 | 1 | 0 | 1 | 2 |
| Percent CNVs with detox genes | 1 | 1.5 | 1.4 | 1.9 | 0.3 | 0 | 0.2 | 0.7 |
|  | GHgam<br>(12) | GM<br>(65) | GNcol<br>(4) | GNgam<br>(38) | GQgam<br>(9) | GW<br>(90) | KE<br>(37) | UGgam<br>(112) |
| CNVs without detox genes | 114 | 388 | 26 | 251 | 111 | 375 | 276 | 336 |
| CNVs with detox genes | 0 | 1 | 0 | 3 | 0 | 1 | 1 | 6 |
| Percent CNVs with detox genes | 0 | 0.3 | 0 | 1.2 | 0 | 0.3 | 0.4 | 1.8 |

Analysis of Gene Ontology (GO) terms for genes encompassed by CNVs showed enrichment for cytochrome P450s. Genes found in CNVs were enriched for 13 molecular function GO terms after multiple correction to a *Q*-value threshold of 0.05 (Supplementary Data S3), primarily reflecting an enrichment for two classes of genes: cytochrome P450s (significant GO terms included monoxygenase activity, heme binding, iron ion binding, oxidoreductase activity) and proteases (significant GO terms included several forms of peptidase activity). No GO terms from biological process or cellular compartment ontologies were significantly enriched.

The 28 detox genes found inside CNVs were predominantly ones that have previously been implicated in insecticide resistance (Table 2, Supplementary Data S2), including the Glutathione S-Transferase Epsilon (*Gste*) cluster and several Cytochrome P450s (*Cyp9k1*, *Cyp6m2*, *Cyp6z1,* and *Cyp6p3*). This again indicates that genes involved in metabolic insecticide resistance have been the focus of amplification events. We therefore performed a detailed analysis of the CNVs in the gene clusters containing *Cyp6p3* (chromosome 2R), *Gste2* (chromosome 3R), *Cyp6m2* (chromosome 3R), *Cyp6z1* (chromosome 3R) and *Cyp9k1* (chromosome X) (Fig. 1a). Since the *Cyp6aa1 / Cyp6aa2* genes, which are adjacent to the *Cyp6p* cluster, were also highly represented in the list of amplified genes (Supplementary Data S2), we extended the study region around *Cyp6p3* to include these genes.

**Table 2:** List of cytochrome P450s, glutathione-S-transferases and carboxylesterase genes found in duplicated regions. None were found on chromosome 3L.

| Chromosome 2L |  | Chromosome 2R |  | Chromosome 3R |  | Chromosome X |  |
| --- | --- | --- | --- | --- | --- | --- | --- |
| Gene ID | Annotation | Gene ID | Annotation | Gene ID | Annotation | Gene ID | Annotation |
| AGAP006724 | COEAE3G | AGAP002862 | CYP6AA1 | AGAP008022 | CYP12F1 | AGAP000818 | CYP9K1 |
| AGAP006725 | COEAE3H | AGAP013128 | CYP6AA2 | AGAP008212 | CYP6M2 |  |  |
| AGAP006726 | COEAE5G | AGAP002863 | COEAE6O | AGAP008219 | CYP6Z1 |  |  |
| AGAP006727 | COEAE6G | AGAP002865 | CYP6P3 | AGAP009190 | GSTE8 |  |  |
| AGAP006728 | COEAE7G | AGAP002866 | CYP6P5 | AGAP009191 | GSTE6 |  |  |
|  |  | AGAP002867 | CYP6P4 | AGAP009192 | GSTE5 |  |  |
|  |  | AGAP002868 | CYP6P1 | AGAP009193 | GSTE4 |  |  |
|  |  | AGAP002869 | CYP6P2 | AGAP009194 | GSTE2 |  |  |
|  |  | AGAP002870 | CYP6AD1 | AGAP009195 | GSTE1 |  |  |
|  |  | AGAP004165 | GSTD2 | AGAP009196 | GSTE7 |  |  |
|  |  |  |  | AGAP009197 | GSTE3 |  |  |
|  |  |  |  | AGAP009696 | CYP325C3 |  |  |

**Figure 1:**
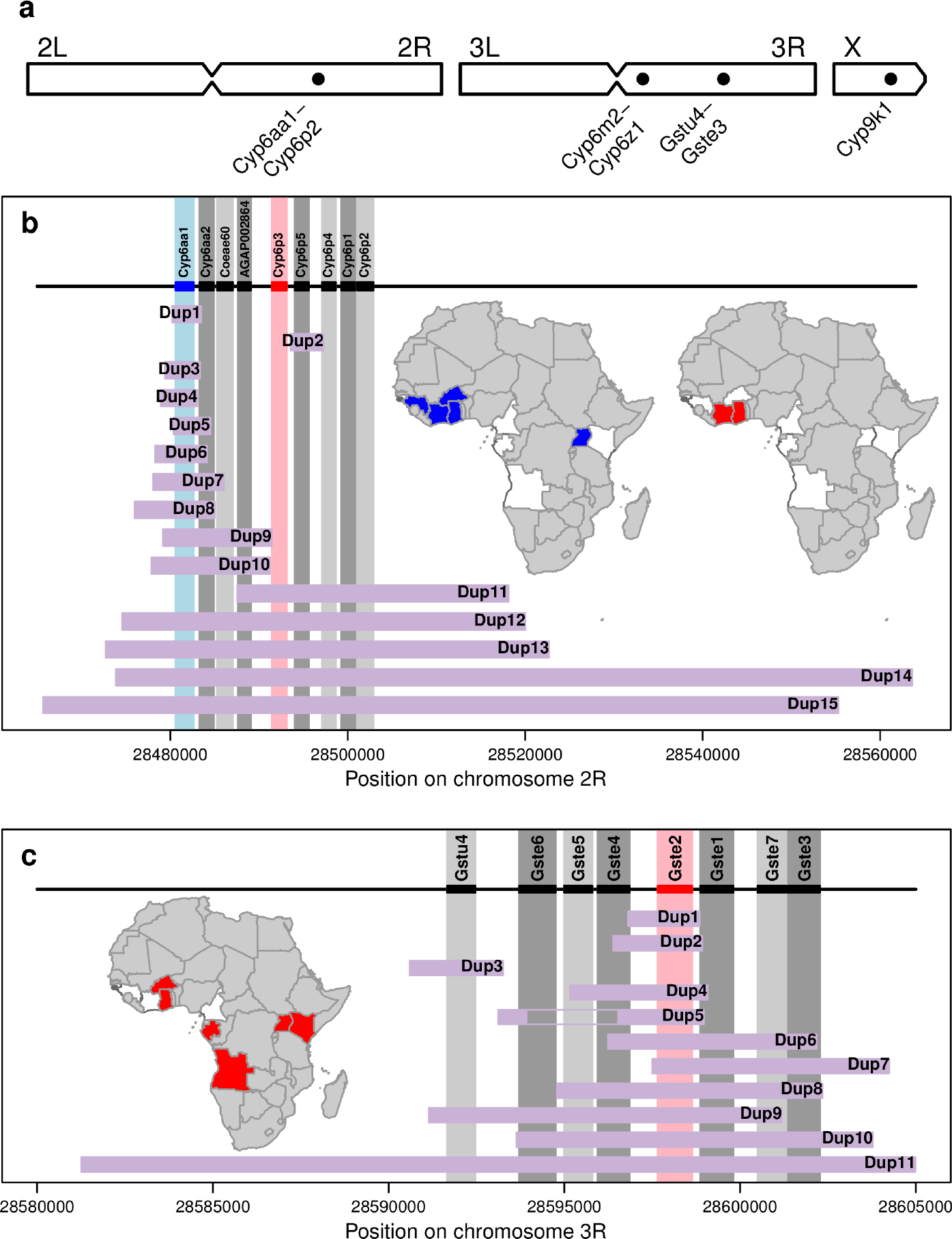
(**a)** CNVs in gene clusters known to be associated with metabolic insecticide resistance were found on all three chromosomes. Inset maps show countries in which at least 5% of individuals carried a CNV in *Cyp6aa1* (blue) and *Cyp6p3* (red), which countries absent from the data set shown in grey. **(b)** Of the 15 CNVs in *Cyp6aa1* - *Cyp6p2*, 13 include *Cyp6aa1* and 5 include *Cyp6p3*. **(c)** Of the 11 duplications in *Gstu4* - *Gste3*, 10 include *Gste2*. Inset map shows countries in which at least 5% of individuals carried a CNV in *Gste2* (red). Black rectangles and vertical grey bars show the positions of the genes in the cluster, with *Cyp6aa1*, *Cyp6p3* and *Gste2* highlighted in colour. Brown horizontal bars show the extent of each CNV, with the gap in Gstue_Dup5 showing the deletion within this amplification. CNV names are abbreviated to Dup* and refer to Cyp6aap_Dup* and Gstue_Dup* in sub-plots **b** and **c** respectively. Further details on each of these CNVs, and of those from the other gene clusters, are presented in Supplementary Data S5-S8.

Using discordant read pairs and reads aligning to CNV breakpoints, we found that three of the five gene clusters showed very high numbers of repeated, independent CNV events. We identified 16 CNV alleles in *Cyp9k1* (named Cyp9k1_Dup1 - 16; Supplementary Data S7 & Supplementary Fig. S4), 15 in the *Cyp6aa1* – *Cyp6p2* cluster (Cyp6aap_Dup1 - 15; Fig. 1b, Supplementary Data S4 & Supplementary Fig. S1), 11 in the *Gstu4* - *Gste3* cluster (Gstue_Dup1 - 11; Fig. 1c, Supplementary Data S5 & Supplementary Fig. S2), one in *Cyp6m2* (Cyp6m_Dup1; Supplementary Data S6 & Supplementary Fig. S3) and one in *Cyp6z3* - *Cyp6z1* (Cyp6z_Dup1; Supplementary Data S6 & Supplementary Fig. S3). Many of these CNVs were found across multiple populations (Supplementary Data S8).

Several CNV alleles were found across different populations (for example, Cyp6aap_Dup7 was found in *An. coluzzii* from Burkina Faso, Côte d’Ivoire, Ghana and Guinea; Supplementary Data S4), although none were found in all populations of either species. Furthermore, multiple CNV alleles covering the same genes could be found in the same population (for example, Cyp9k1_Dup4, Dup11 and Dup15 in *An. gambiae* from Burkina Faso; Supplementary Data S7). This resulted in some genes being amplified at very high frequency in certain populations (Table 3). In particular, over 92% of individuals had a CNV in *Cyp9k1* in *An. gambiae* from Burkina Faso, and 90% of individuals had a CNV covering all genes in the *Cyp6p* cluster in *An. coluzzii* from Côte d’Ivoire.

**Table 3:** Number (and proportion) of individuals with a CNV covering *Cyp6aa1*, *Cyp6p3*, *Gste2*, *Cyp6m2*, *Cyp6z1* or *Cyp9k1*. CNVs were identified using discordant and breakpoint reads. AO = Angola, BF = Burkina Faso, CI = Côte d’Ivoire, CM = Cameroon, FR = French Mayotte, GA = Gabon, GH = Ghana, GM = The Gambia, GN = Guinea, GQ = Equatorial Guinea, GW = Guinea-Bissau, KE = Kenya, UG = Uganda. col = *An. coluzzii*, gam = *An. gambiae*. Numbers in brackets after the population name indicate the total number of samples from that population. Proportions above 50% have been highlighted in bold.

|  | AOcol<br>(78) | BFcol<br>(75) | BFgam<br>(92) | CIcol<br>(71) | CMgam<br>(297) | FRgam<br>(24) | GAgam<br>(69) | GHcol<br>(55) |
| --- | --- | --- | --- | --- | --- | --- | --- | --- |
| Cyp6aa1 | 0 | **68 (90.7%)** | 3 (3.3%) | **63 (88.7%)** | 2 (0.7%) | 0 | 0 | 6 (10.9%) |
| Cyp6p3 | 0 | 3 (4%) | 0 | **64 (90.1%)** | 0 | 0 | 0 | 3 (5.5%) |
| Gste2 | 11 (14.1%) | 18 (24%) | 0 | 0 | 13 (4.4%) | 0 | 9 (13%) | 6 (10.9%) |
| Cyp6m2 | 0 | 0 | 0 | 5 (7%) | 0 | 0 | 0 | 0 |
| Cyp6z1 | 0 | 0 | 15 (16.3%) | 0 | 0 | 0 | 0 | 0 |
| Cyp9k1 | 12 (15.4%) | 1 (1.3%) | **85 (92.4%)** | 0 | 14 (4.7%) | 0 | 0 | 3 (5.5%) |
|  | GHgam<br>(12) | GM<br>(65) | GNcol<br>(4) | GNgam<br>(40) | GQgam<br>(9) | GW<br>(91) | KE<br>(48) | UGgam<br>(112) |
| Cyp6aa1 | 0 | 0 | **3 (75%)** | 1 (2.5%) | 0 | 0 | 0 | **72 (64.3%)** |
| Cyp6p3 | 0 | 0 | 0 | 0 | 0 | 0 | 0 | 1 (0.9%) |
| Gste2 | 0 | 0 | 0 | 0 | 0 | 0 | 23 (47.9%) | 19 (17%) |
| Cyp6m2 | 0 | 0 | 0 | 0 | 0 | 0 | 0 | 0 |
| Cyp6z1 | 0 | 0 | 0 | 5 (12.5%) | 0 | 0 | 0 | 0 |
| Cyp9k1 | **6 (50%)** | **38 (58.5%)** | 0 | **34 (85%)** | 0 | 10 (11%) | 0 | 37 (33%) |

### 2.2 CNVs in metabolic resistance genes are under positive selection

Several CNV alleles were found at high local frequencies (Supplementary Data S8), suggesting that they are likely to be under positive selection. To investigate this possibility, we phased the CNV genotype calls onto the Ag1000G phase 2 haplotype scaffold and calculated extended haplotype heterozygosity (EHH) for the CNV alleles present in at least 5% of individuals in a population.

Rates of EHH decay around CNV alleles were consistently lower than for wild-type haplotypes (Fig. 2, S5, S7 and S9), supporting our contention that these alleles are reaching high frequency through positive selection. Furthermore, the median length of shared haplotypes was significantly higher between pairs of haplotypes carrying the same CNV allele than between wild-type haplotypes from the same population (bootstrapped 95% confidence intervals for the medians did not overlap, Fig. 3, S6, S8 and S10).

**Figure 2:**
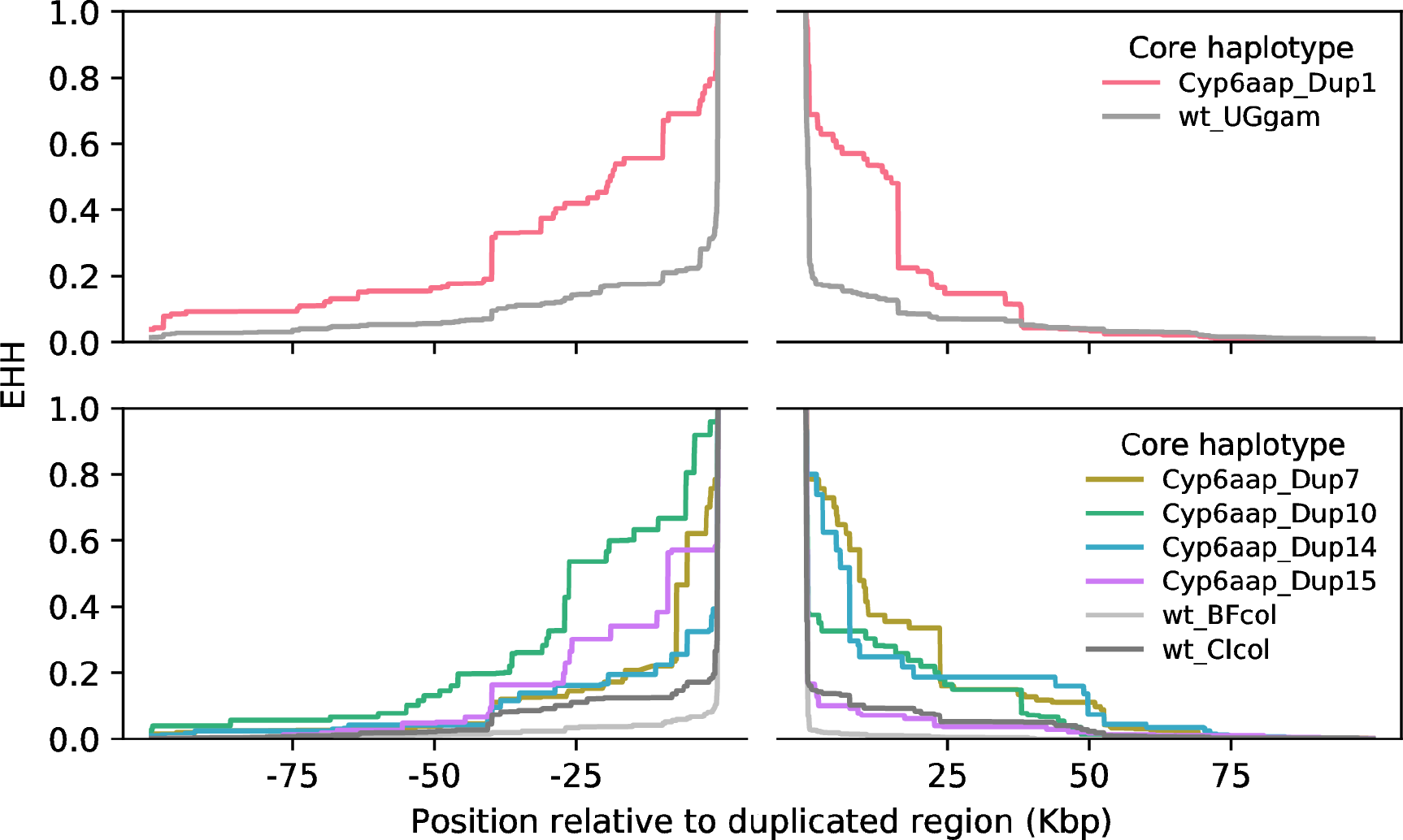
Evidence for prolonged linkage disequilibrium around CNVs in the *Cyp6aa1-Cyp6p2* gene cluster. Extended Haplotype Heterozygosity (EHH) decay was calculated around CNV and non-CNV (wt) haplotypes using SNPs from outside the region containing CNVs (break in the x axis). BF = Burkina Faso, CI = Côte d’Ivoire, UG = Uganda, col = *An. coluzzii*, gam = *An. gambiae*.

**Figure 3:**
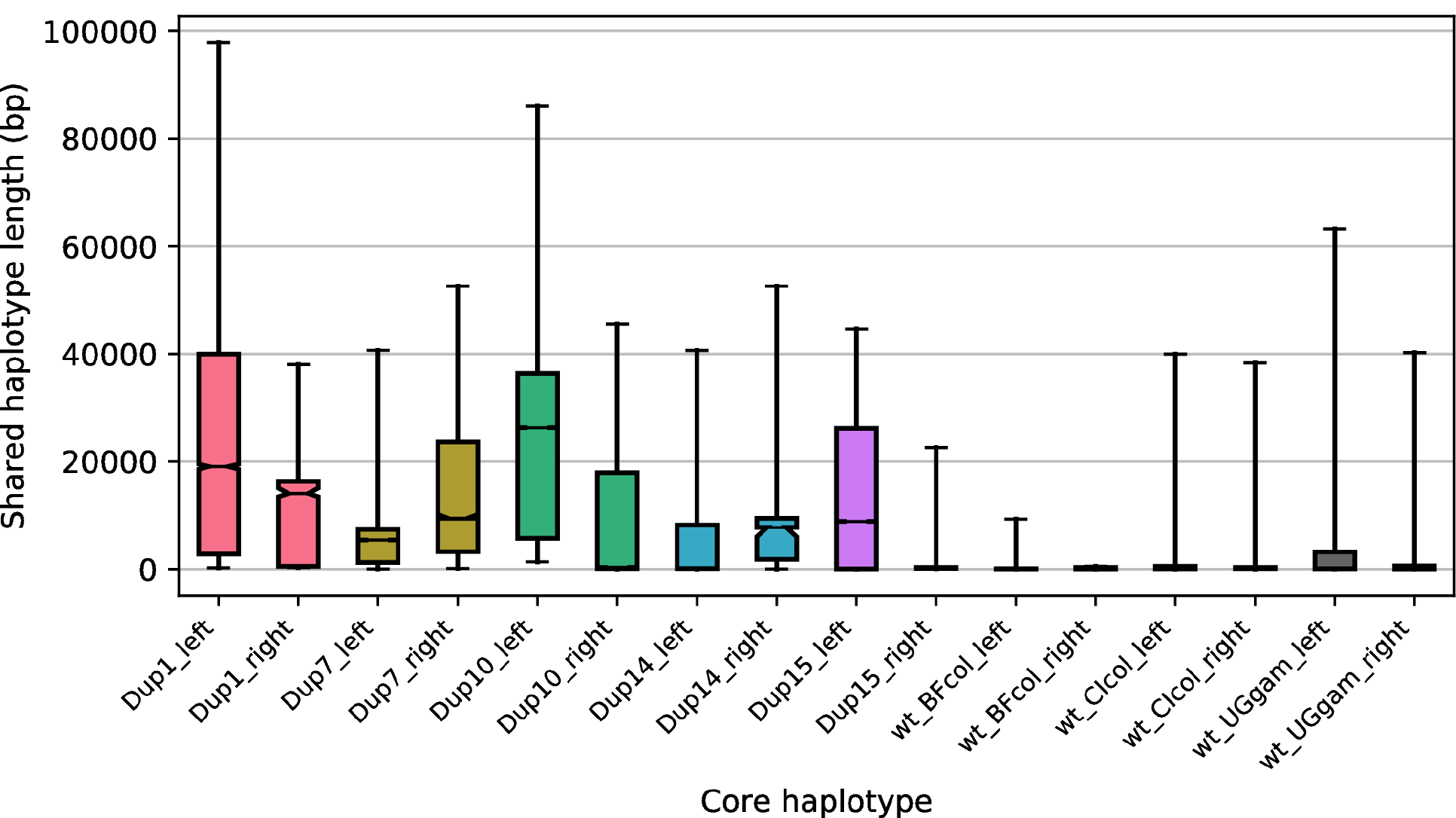
Lengths of pairwise shared haplotypes are greater between samples sharing a CNV allele than between wild-type samples. Shared haplotype lengths were calculated on either side of the CNV-containing region of the *Cyp6aa/p* gene cluster. Non-CNV (wt) samples were taken from the same populations as the focal CNV alleles. Bars show the distribution of shared haplotype lengths between all haplotype pairs with the same core haplotype. Bar limits show the inter-quartile range, fliers show the 5th and 95th percentiles, horizontal black lines show the median, notches in the bars show the bootstrapped 95% confidence interval for the median. The names of the CNVs (Cyp6aap_Dup*) are abbreviated as Dup*. BF = Burkina Faso, CI = Côte d’Ivoire, UG = Uganda, col = *An. coluzzii*, gam = *An. gambiae*.

Phasing of CNV genotype calls was only possible for simple duplications, where the zygosity of the CNV alleles could be determined from the copy number estimates. For CNV alleles with higher copy numbers (triplications and above), this was not possible, and thus the EHH decay could not be calculated. In the case of *Cyp9k1*, the CNV with the highest frequency (Cyp9k1_Dup11, found in *An. gambiae* from Burkina Faso, Ghana and Guinea) could not be phased. We therefore investigated whether this CNV was associated, at the sample level, with haplotypes under selection. Hierarchical clustering of the haplotypes in these three populations revealed two large cross-population haplotype clusters around *Cyp9k1*, indicating selective sweeps (Fig. S11). Cluster 1 was very strongly associated with Cyp9k1_Dup11 in both males (Fisher’s exact test, *P* < 0.0001; Supplementary Table S1) and females (Spearman’s rank correlation: ρ = 0.9, *P* < 0.0001, Fig. 4a). Cluster 2 was associated with the presence of Cyp9k1_Dup15, but the correlation was not as close as between Cluster 1 and Cyp9k1_Dup11 (Spearman’s rank correlation: ρ = 0.65, *P* < 0.0001, Fig. 4b & Supplementary Table S2).

**Figure 4:**
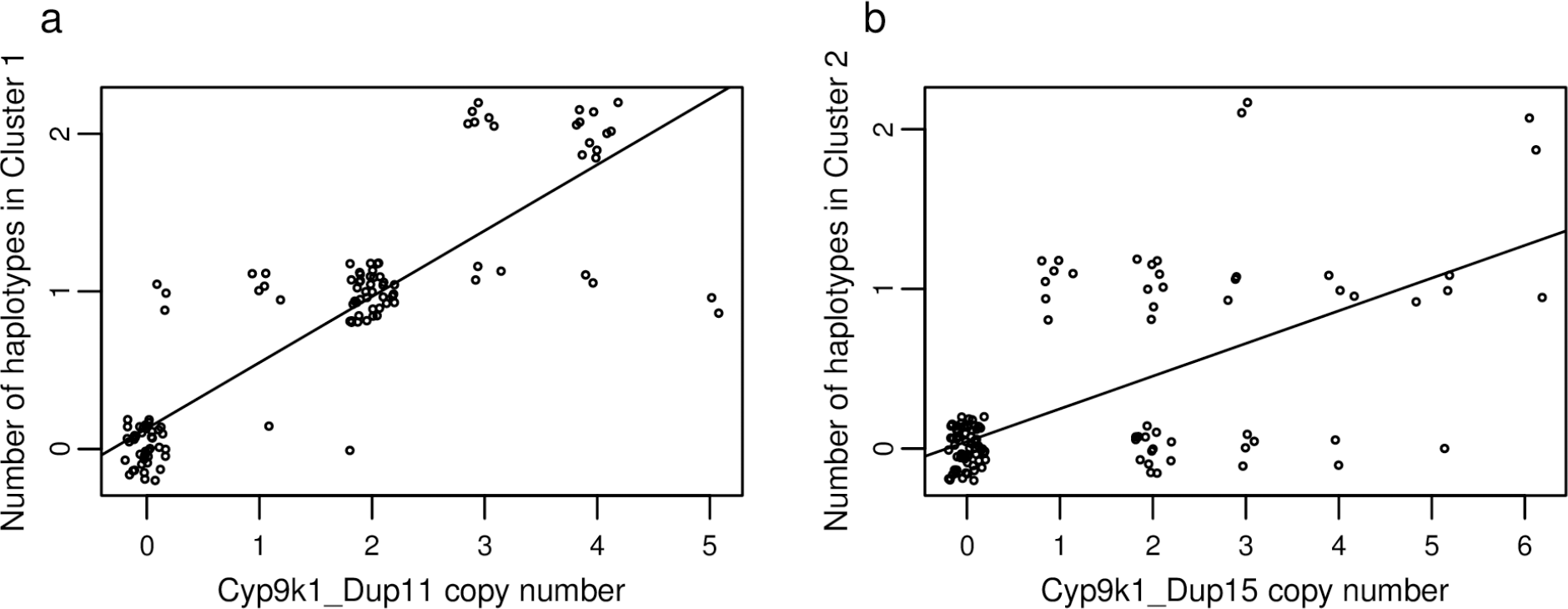
The two main haplotype clusters around *Cyp9k1* in Burkina Faso, Ghana and Guinea are associated with respective CNV alleles. Points are jittered to show overlapping data. Lines show least squares regression through the data. (a) Strong correlation between Cyp9K1_Dup11 and haplotype Cluster 1. Most of the points lie on a line of slope 0.5, indicating that Cyp9k1_Dup11 is found most frequently as a triplication (two extra copies per chromosome), although both lower and higher copy number versions of this CNV exist. (b) Weaker correlation between Cyp9k1_Dup15 and haplotype Cluster 2.

### 2.3 A *Gste2* duplication in Burkina Faso is associated with the resistance-conferring I114T mutation

Ten of the 11 CNV alleles found in the *Gstu4* - *Gste3* cluster included *Gste2* (Fig. 1), perhaps reflecting the known importance of this gene in insecticide resistance. The well-characterised I114T mutation in *Gste2* is known to confer DDT resistance [28], and could be associated with gene duplications in a similar fashion to other mutations such as the *Ace-1* G119S. We therefore investigated whether any of the CNV alleles in *Gste2* were associated with this mutation. 114T is present across Africa and in both *An. gambiae* an *An. coluzzii* [36], but was only associated with Gstue_Dup1 in our data. Gstue_Dup1 was found in 16 *An. coluzzii* samples from Burkina Faso, all of which were at least heterozygote for 114T (Supplementary Table S3). The presence of 114T homozygotes, together with the ratio of reads supporting the I114 and 114T alleles in heterozygotes (roughly 1:2), indicate that both copies of *Gste2* in the Gstue_Dup1 CNV carry the 114T mutation.

### 2.4 Cyp6aa1 is more strongly associated with CNVs than Cyp6p3

Of the 15 CNV alleles found in the *Cyp6aa1* – *Cyp6p2* cluster, five included *Cyp6p3* but 13 included *Cyp6aa1* (Supplementary Data S4). *Cyp6p3* CNVs were found at high (> 50%) frequency in one population (Côte d’Ivoire *An. coluzzii*: 90%), while *Cyp6aa1* CNVs were found at high frequency in *An. coluzzii* from Burkina Faso (91%), Côte d’Ivoire (89%) and Guinea (75%), and in *An. gambiae* from Uganda (64%).

## 3 Discussion

Our study detected 1557 CNVs in *An. gambiae* and *An. coluzzii* and revealed a striking enrichment for gene families involved in metabolic insecticide resistance. These results mirror findings in *Drosophila melanogaster*, where cytochrome P450s were disproportionately represented in CNVs [37]. Similarly, in *Aedes aegypti*, cytochrome P450s were enriched among genes showing evidence of higher copy number in populations resistant to deltamethrin compared to susceptible populations [38]. Strikingly, the five metabolic genes most strongly associated with insecticide resistance in the literature for *An. gambiae* and *An. coluzzii*, and which have been shown to metabolise insecticides in vitro (*Gste2*, *Cyp6p3*, *Cyp6m2*, *Cyp6z1* and *Cyp9k1*), were all found to be amplified in at least one population in our dataset. Furthermore, three of these genes showed evidence of repeated independent CNV events within and between populations, with as many as 16 independent CNV alleles in *Cyp9k1*.

Expression of *Gste2* is higher in DDT resistant *An. gambiae* [39,40] and *An. funestus* [41] compared to susceptible mosquitoes, and transgenic expression of *An. gambiae* / *An. funestus Gste2* in *Drosophila* provides resistance to DDT [28,41]. Non-synonymous SNPs in *Gste2* have also been shown to be associated with resistance to DDT in both *An. gambiae* [28] and *An. funestus* [41]. In our study, *Gste2* was amplified in Kenya, in *An. coluzzii* from Angola, Burkina Faso and Ghana, and in *An. gambiae* from Gabon and Uganda.

*Cyp6p3* is up-regulated in mosquitoes resistant to pyrethroids, DDT and bendiocarb [29,30,42-45], metabolises permethrin and deltamethrin [30] and provides resistance to pyrethroids when expressed in *Drosophila* [29]. *Cyp6m2* is also up-regulated in mosquitoes with resistance to permethrin, DDT and bendiocarb [29,31,42,46], metabolises pyrethroids and DDT [31,32] and provides resistance to pyrethroids, DDT and bendiocarb when expressed in *Drosophila* [29]. In our study, *Cyp6p3* and *Cyp6m2* were found amplified primarily in *An. coluzzii* from Côte d’Ivoire. Interestingly, mosquitoes from the Côte d’Ivoire population that our samples were drawn from over-express both *Cyp6p3* and *Cyp6m2* compared to susceptible populations [29]. Particularly in the case of *Cyp6m2*, this over-expression is unlikely to be driven solely by CNVs, since the CNV frequency and copy number are not sufficient to explain the expression levels, but the selective pressure to up-regulate these genes may have played a part in maintaining these CNVs in the population.

*Cyp6z1* was amplified in *An gambiae* from Burkina Faso and Guinea. *Cyp6z1* is up-regulated in mosquitoes with resistance to pyrethroids and DDT [39,47] and metabolises DDT and carbaryl [33]. Finally, *Cyp9k1* was the most widely amplified gene of the five, with CNVs found in over half of the populations in our dataset. *Cyp9k1* is up-regulated in mosquitoes resistant to pyrethroids and DDT [44,45,48], and metabolises deltamethrin [35]. Furthermore, a selective sweep in the *Cyp9k1* region has been associated with insecticide resistance in *An. coluzzii* [34].

In-depth investigation of the CNVs around these five genes revealed strong evidence that they provide a selective advantage. First, some of the CNV alleles were found at high frequencies and across several populations. Second, the CNV alleles consistently showed evidence of being under positive selection as haplotype homozygosity was extended further for the CNVs than for wild-type haplotypes. Evidence for positive selection was also found in a CNV where the EHH score could not be calculated. Cyp9k1_Dup11, which exists as both duplications and triplications and thus could not be phased onto a haplotype scaffold for homozygosity calculation, was consistently found in the same samples as the haplotype of a large selective sweep around *Cyp9k1* in *An. gambiae* from Burkina Faso, Guinea and Ghana, raising the strong possibility that this CNV is the focus of the selective sweep. It cannot be excluded that Cyp9k1_Dup11 is in linkage disequilibrium with another mutation that is driving the sweep. However, the high frequency of the triplicated version of Cyp9k1_Dup11 compared to the duplicated version, both of which are associated with the swept haplotype cluster, suggests that higher-order amplifications of *Cyp9k1* provide a selective advantage. The changes in allele frequencies in the different amplification levels in this CNV will need to be monitored to determine whether the triplication eventually replaces the duplication entirely.

We found multiple independent CNVs in three of the five gene regions, with 11 CNV alleles around *Gste2* (a gene cluster containing *Gste1-7* and *Gstu4*), 15 around *Cyp6p3* (a gene cluster including *Cyp6aa1-2* and *Cyp6p1-5*) and 16 around *Cyp9k1*. For the *Cyp6aa/p* cluster, these independent CNVs were primarily found in *An. coluzzii* from Burkina Faso, Côte d’Ivoire and Ghana. In *Cyp9k1*, CNVs were primarily found in *An. gambiae* from Burkina Faso and Ghana and Guinea.

All but one of the eleven CNV alleles in the *Gstu/e* cluster included *Gste2*, indicating that this is the major target of gene amplification in this cluster. Given the body of evidence linking *Gste2* to DDT and pyrethroid resistance across multiple species (*An. gambiae*: [28], *An. funestus*: [41], *Aedes aegypti*: [49]), the focus of amplifications on this gene is likely to be linked to its importance in resistance.

Interestingly, the Gstue_Dup1 duplication in Burkina Faso occurs on the background of the *Gste2*_114T SNP, associated with DDT resistance in *An. gambiae* [28]. The duplication may therefore serve to increase the dosage of *Gste2*, whose detoxifying activity has already been elevated by the 114T mutation. Alternatively, the role of Gstue_Dup1 may be to compensate for any negative fitness effects of 114T. Whilst impaired *Gste2* activity may be compensated by increasing the expression of the gene, Gstue_Dup1 is homogeneous for 114T, excluding the possibility of compensation by pairing of mutant and wild-type alleles as found in heterogeneous *Ace-1* duplications [6].

Unexpectedly, in the *Cyp6aa/P* cluster, only five of the 15 CNVs included *Cyp6p3*, and these were only found at appreciable frequency in *An. coluzzii* from Côte d’Ivoire. In contrast, 13 of the 15 CNVs included *Cyp6aa1*, with high CNV frequencies found in *An. coluzzii* from Burkina Faso, Côte d’Ivoire and Guinea, and in *An. gambiae* from Uganda. Furthermore, the five high frequency CNVs that include *Cyp6aa1* (Cyp6aap_Dup1, Cyp6aap_Dup7, Cyp6aap_Dup10, Cyp6aap_Dup14, Cyp6aap_Dup15) all show evidence of positive selection. While *Cyp6aa1* has received substantially less attention than *Cyp6p3*, it has previously been implicated in insecticide resistance. Expression of *Cyp6aa1* is higher in populations of *An. gambiae* and *An. coluzzii* that are resistance to pyrethroids and DDT compared to susceptible laboratory colonies [43,48]. There is also strong evidence for a link between *Cyp6aa1* and insecticide resistance in two congeneric species. In *An. funestus* expression of *Cyp6aa1* is higher in mosquitoes that have survived permethrin exposure compared to a susceptible strain [50,51], and the protein has been shown to metabolise pyrethroids and drive resistance when expressed in *Drosophila* [50]. In *An. minimus*, the ortholog of *Cyp6aa1* is up-regulated as a result of selection for resistance to deltamethrin [52], and the protein has been shown to metabolise pyrethroids [53]. The ability of *An. gambiae Cyp6aa1* to metabolise insecticides has not been tested empirically, although theoretical modelling suggests that it should effectively bind to permethrin and deltamethrin [50]. The high frequency of amplifications in *Cyp6aa1* and the signals of selection associated with them suggest that the importance of this gene for insecticide resistance in *An. gambiae* and *An. coluzzii* has been under-appreciated.

In conclusion, our results show a key role for CNVs in the adaptive response to strong and evolutionary recent selective pressure. In populations of *Anopheles* mosquitoes across Africa, genes involved in metabolic resistance to insecticides have been duplicated and these duplications have been driven to high frequencies by positive selection. These results highlight CNVs as a form of variation that can act as a front-line response to selective pressures requiring changes in expression levels, perhaps because low-copy-number whole-gene amplifications have relatively little negative effect on fitness and can thus exist in the standing genetic variation of a population without being removed by purifying selection. Our findings also highlight *Cyp6aa1* as a gene that should be more closely investigated for its importance in *An. gambiae*, having been so far overlooked in preference to its genomic neighbour *Cyp6p3*. More broadly, the focus on SNPs in *An. gambiae* insecticide resistance research has allowed the emergence and selective spread of copy number mutations in key insecticide resistance genes to go unnoticed. Our findings demonstrate the importance of surveillance and investigation of CNVs in these genes. To this end, the breakpoint descriptions provided in our study will allow these CNVs to be screened and monitored in mosquito populations, allowing the spread of these mutations to be tracked and providing the groundwork for future studies investigating their resistance profile.

## 4 Methods

### 4.1 Population sampling and whole genome sequencing

We analysed data from 1,142 individual wild-caught specimens of *An. gambiae* and *An. coluzzii* collected and sequenced in phase 2 of Ag1000G [13]. The specimens were collected from sites in 13 African countries (Angola *An. coluzzii* n = 78, Burkina Faso *An. coluzzii* n = 75, Burkina Faso *An. gambiae* n=92, Cameroon *An. gambiae* n = 297, Côte d’Ivoire *An. coluzzii* n = 71, Equatorial Guinea (Bioko) *An. gambiae* n = 9, Gabon *An. gambiae* n = 69, Ghana *An. coluzzii* n = 55; Ghana *An. gambiae* n = 12, Guinea *An. coluzzii* n = 4, Guinea *An. gambiae* n = 40, Guinea-Bissau (mixed ancestry) n = 91, Kenya (undetermined ancestry) n = 48, Mayotte *An. gambiae* n = 24, The Gambia (mixed ancestry) n = 65, Uganda *An. gambiae* n = 112). Individual specimens were sequenced using the Illumina Hi-Seq platform to obtain 100 bp paired-end reads with a target coverage of 30X. Further details of population sampling, sample preparation, sequencing, alignment, species identification and data production are reported elsewhere [36]

### 4.2 Calculation and normalisation of coverage

For each individual, we used the *pysam* software package (https://github.com/pysam-developers/pysam) to count the number of aligned reads (coverage) in non-overlapping 300 bp windows over the nuclear genome. The position of each read was considered to be its alignment start point, thus each read was only counted once. Sequencing coverage can be biased by variation in local nucleotide composition. To account for this, we computed a normalised coverage from the read counts based on the expected coverage of each window given its GC content [54]. For each 300 bp window we computed the percentage of (G+C) nucleotides to the nearest percentage point within the reference sequence and then divided the read counts in each window by the mean read count over all autosomal windows with the same (G+C) percentage. To minimise the impact of copy number variation when calculating these normalising constants, we excluded windows from the calculation of mean read counts where previous analyses of genome accessibility have found evidence for excessively high or low coverage or ambiguous alignment (windows with <90% accessible bases according to the Ag1000G phase 2 genome accessibility map [13], referred to as “accessible windows”). The normalised coverage values were then multiplied by a factor of 2, so that genome regions with a normal diploid copy number should have an expected normalised coverage of 2.

Before examining the normalised coverage data for evidence of copy number variation, we applied two filters to exclude windows where coverage may be an unreliable indicator of copy number. The first filter removed windows where more than 2% of reads were aligned with mapping quality 0 (Fig. S12), which indicates that a read is mapped ambiguously and could be mapped equally well to a different genomic location. This filter removed 159,587 (20.8%) of 768,225 windows. The second filter removed windows where the percentage (G+C) content was extreme and rarely represented within the accessible reference sequence (fewer than 100 accessible windows with the same (G+C) percentage) because the small number of windows makes the calculation of a (G+C) normalising constant unreliable. This filter removed 13,484 (2.2%) of the 608,638 remaining windows. Windows retained for analysis were referred to as “filtered windows”.

### 4.3 Genome-wide copy number variation discovery

To detect the most likely copy-number state (CNS) at each window in each individual, we applied a Gaussian Hidden Markov Model (HMM) to the individual’s normalised windowed coverage data, following a similar approach to [55] and [5] (see Supplementary Methods SM1 for details). Since we are primarily interested in amplifications rather than deletions, we obtained a raw set of CNV calls for each sample by locating contiguous runs of at least five windows with amplified CNS (CNS > 2, or CNS > 1 for chromosome X in males).

### 4.4 CNV filtering

From the raw CNV call set, we created a quality-filtered list of CNV calls. We first removed samples with very high coverage variance, since high variance could lead to erratic CNV calls. We therefore removed 27 samples where the variance in normalised coverage was greater than 0.2 (Fig. S13), retaining 1,115 samples for further analysis.

We then applied two filters to the raw CNV calls from these 1,115 samples. For the first filter, we computed likelihoods for each raw CNV call for both the copy number state predicted by the HMM and for a null model of copy number = 2, and removed CNV calls where the likelihood ratio was < 1000 (Supplementary Methods SM2). For the second filter, we removed CNVs with low population frequencies. To do this, the raw CNV calls needed to be matched so that the same CNV in different individuals could be identified. We classed any two CNVs as identical if the breakpoints predicted by their copy number state transitions were within one window of each-other. We then removed CNVs that were not found in at least 5% of individuals in at least one population (or at least 3 individuals for populations smaller than 40).

### 4.5 Discovery of gene duplications and gene enrichment analysis

To determine the genes contained within each CNV, we compared the start and end points of the CNVs to the start and end points of all genes listed in the AgamP4.2 gene annotations (*Anopheles-gambiae-PEST_BASEFEATURES_AgamP4.2.gff3*). The start / end points of each CNV were calculated as the median of the start / end points of all the raw CNV calls that were matched to it. To keep only genes for which good coverage data were available, we retained only genes containing at least 50% filtered windows. We classed a retained gene as copied by a CNV if all of the filtered windows within the gene were inside the CNV. We performed simulations to determine whether the CNVs that we detected contained more genes than expected by chance. For each run of the simulation, we randomly re-allocated the start positions of every detected CNV, keeping the number of filtered windows covered by the CNVs unchanged, and calculated the number of CNVs that included at least one gene. This simulation was run 10,000 times to obtain the distribution of the null model.

We identified genes that could potentially be involved in metabolic resistance through detoxification (“metabolic detox genes”) by finding genes whose annotations contained the terms “P450”, “glutathione S-transferase” or “carboxylesterase” in the AgamP4 transcript annotations (*Anopheles-gambiae-PEST_TRANSCRIPTS_AgamP4.2.fa*). We performed simulations to determine whether genes copied by CNVs were enriched for detox genes. For each run of the simulation, we randomly re-allocated the genes encompassed by each CNV, keeping the number of consecutive genes covered by each CNV unchanged, and calculated the number of CNVs that included at least one metabolic detox gene. This simulation was run 10,000 times to obtain the distribution of the null model.

GO term analysis of genes included in CNVs was performed using the *topGO* [56] package in R. False discovery rates were calculated from the *P*-values using the R package *fdrtool* [57].

### 4.6 Identifying CNV alleles at candidate metabolic insecticide resistance loci

We characterised in detail the independent duplication events (CNV alleles) at five gene clusters of particular interest (*Cyp6aa1* – *Cyp6p2*, *Gstu4* – *Gste3*, *Cyp6m2* – *Cyp6m4*, *Cyp6z3* – *Cyp6z1*, *Cyp9k1*) using their unique patterns of discordant read pairs and reads crossing the CNV breakpoint (breakpoints reads, see Supplementary Methods SM3 and Fig. 5). We manually inspected the five regions of interest in all 1142 samples to identify patterns of discordant and breakpoint reads (“diagnostic reads”) consistently associated with changes in coverage (Supplementary Fig. S1-S4). The start and end point of each CNV allele could usually be precisely determined by the breakpoint reads, and was otherwise determined by discordant read pairs or the point of change in coverage (Supplementary Data S4-S7). Once the diagnostic reads were identified for a CNV allele, we recorded the presence of that allele in all samples with at least two supporting diagnostic reads.

**Figure 5:**
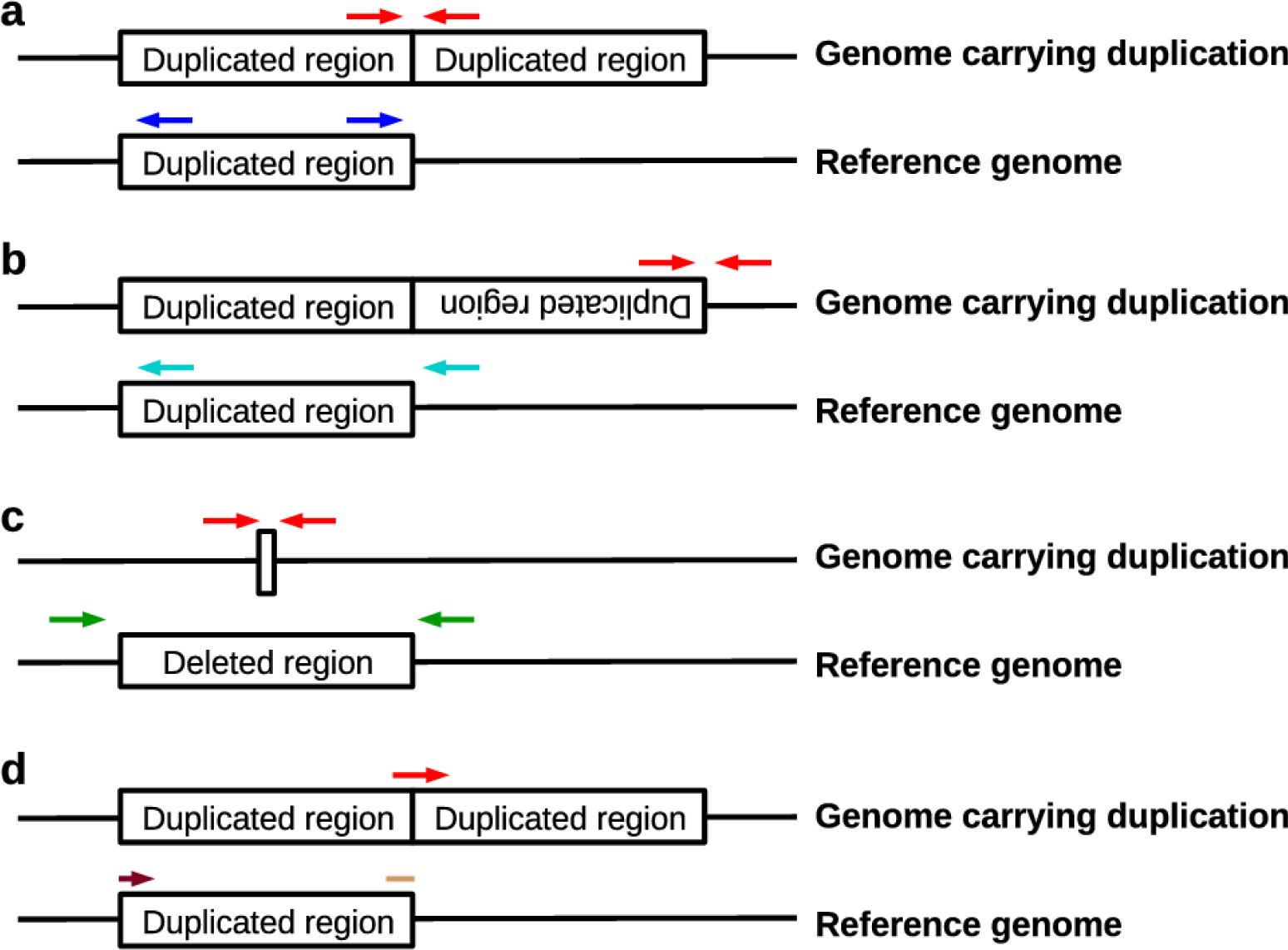
Three types of discordant read pairs (**a**-**c**) and breakpoint reads (**d**) were used to identify different CNV alleles. **a** In tandem duplications, read pairs derived from segments spanning the CNV breakpoint (red arrows) align facing away from each other around the breakpoint on the reference genome (dark blue arrows). **b** In tandem inversions, read pairs derived from segments spanning the end of the inverted segment (red arrows) align facing in the same direction as each other around the breakpoint on the reference genome (cyan arrows). **c** In deletions, read pairs derived from segments spanning the deleted sequence (red arrows) align in the correct orientation around the breakpoint, but farther apart than expected given the insert size of the sequencing library (green arrows). **d** In any of the above types of CNV (tandem duplication shown here as an example), reads crossing the breakpoint (red arrow) will only partially align on either side of the breakpoint. For the tandem duplication shown here, the start of the read (light brown start of an arrow) aligns at the end of the duplicated region, while the end of the read (dark brown end of an arrow) aligns at the start of the duplication.

### 4.7 Detecting signals of selection on CNV alleles

We used the phased haplotypes to calculate the pairwise shared haplotype length and the EHH for each CNV allele [58], using only SNPs from outside the region in which CNVs were found. EHH calculations were performed using the *scikit-allel* Python package [ref].

Haplotypes clusters in the *Cyp9k1* region were were obtained with *scikit-allel*, using the first 1000 SNPs on the centromeric side of *Cyp9k1* (the telomeric side of this gene has low levels of accessibility). A distance matrix between haplotypes was computed using the proportion of accessible SNPs that differed between pairwise haplotype combinations. This was used to perform hierarchical clustering, with haplotype clusters defined using a cut-off of 0.001.

### 4.8 Statistics

Statistical analysis was performed in R [59]. Contingency tables were analysed with Fisher’s exact test. Where the sample size was too large for the Fisher’s exact test, *P* values were obtained using the “simulated.p.value” option with 10^6^ replicates.

### 4.9 Estimating allele-specific copy numbers and phasing CNV genotypes onto the Ag1000G haplotype scaffolds

To determine the allele-specific copy number in a sample, we estimated the change in coverage associated with each CNV allele (Supplementary Methods SM4 and Fig. S14). Thus, even when overlapping CNV alleles were present in a single sample, we were usually able to determine the number of copies of each allele.

For single-copy CNVs, it is possible to determine the genotype of a sample from the copy numbers (copy numbers of 1 indicate a heterozygote, copy numbers of 2 indicate a homozygote for the CNV). For higher-order CNVs, this is not possible because a heterozygote triplication cannot be distinguished from a homozygote duplication. We therefore applied two filters to retain only single-copy CNV alleles. The first filter removed CNV alleles where the allele-specific copy number was found to rise above 2 in the data (if only a single sample rose as high as 2.5, we assumed that this could be an error and classed it as 2). This filter removed five CNV alleles (Cyp6aap_Dup11, Gstue_Dup2, Gstue_Dup8, Cyp9k1_Dup11, Cyp9k1_Dup15). For the second filter, we classed each sample as homozygote wild-type, heterozygote or homozygote CNV based on their copy numbers, and then removed CNV alleles that were inconsistent with Hardy-Weinberg expectations within the populations in which they were found. This filter removed four CNV alleles (Cyp6aap_Dup4, Gstue_Dup5. Gstue_Dup7, Cyp9k1_Dup10). Three CNV alleles (Cyp9k1_Dup7, Cyp9k1_Dup13 and Cyp9k1_Dup14) were also excluded because of difficulties in calling allele-specific copy number (Supplementary Data S7). In one case (Cyp6m2_Dup1), all individuals were found to have a copy number of 2, indicating that the CNV is a triplication, with no duplications present in the population. This CNV was therefore retained, with all samples carrying the CNV classed as heterozygote.

CNV alleles that passed both filters were phased onto the Ag1000G phase 2 haplotype scaffolds using the *MVNCALL* software v1.0 [60], using default parameters apart from setting λ = 0.1 to ensure that none of the input CNV genotype calls were changed during phasing. For each of the five gene clusters, phasing was performed using the 200 non-singleton SNPs either side of the region in which CNVs were found, thus avoiding the inclusion of SNPs found inside any of the CNVs. Haplotypes that contained more than one CNV allele were rare and therefore excluded from subsequent calculations of haplotype heterozygosity.

## Acknowledgements

This work was supported by the Wellcome Trust (090770/Z/09/Z; 090532/Z/09/Z; 098051), Medical Research Council UK (MR/P02520X/1; MR/M006212/1) and the National Institute of Allergy and Infectious Diseases (NIAID. R01-AI116811). The content is solely the responsibility of the authors and does not necessarily represent the official views of the NIAID or NIH.

## References

[1] Chen Z, Cheng C-HC, Zhang J, Cao L, Chen L, Zhou L, et al. Transcriptomic and genomic evolution under constant cold in Antarctic notothenioid fish. Proceedings of the National Academy of Sciences. 2008;105: 12944–12949.

[2] Emerson JJ, Cardoso-Moreira M, Borevitz JO, Long M. Natural selection shapes genome-wide patterns of copy-number polymorphism in *Drosophila melanogaster*. Science. 2008;320: 1629– 1631.

[3] Redon R, Ishikawa S, Fitch KR, Feuk L, Perry GH, Andrews TD, et al. Global variation in copy number in the human genome. Nature. 2006;444: 444–454.

[4] Handsaker RE, Van Doren V, Berman JR, Genovese G, Kashin S, Boettger LM, et al. Large multiallelic copy number variations in humans. Nature Genetics. 2015;47: 296.

[5] Leffler EM, Band G, Busby GBJ, Kivinen K, Le QS, Clarke GM, et al. Resistance to malaria through structural variation of red blood cell invasion receptors. Science. 2017;356: eaam6393.

[6] Assogba BS, Djogbénou LS, Milesi P, Berthomieu A, Perez J, Ayala D, et al. An *ace-1* gene duplication resorbs the fitness cost associated with resistance in *Anopheles gambiae*, the main malaria mosquito. Scientific Reports. 2015;5: 14529.

[7] Labbé P, Berticat C, Berthomieu A, Unal S, Bernard C, Weill M, et al. Forty years of erratic insecticide resistance evolution in the mosquito *Culex pipiens*. PLoS Genetics. 2007;3: e205.

[8] Sudmant PH, Mallick S, Nelson BJ, Hormozdiari F, Krumm N, Huddleston J, et al. Global diversity, population stratification, and selection of human copy-number variation. Science. 2015;349.

[9] Kiszewski A, Mellinger A, Spielman A, Malaney P, Sachs SE, Sachs J. A global index representing the stability of malaria transmission. The American Journal of Tropical Medicine and Hygiene. 2004;70: 486–498.

[10] Bass C, Field LM. Gene amplification and insecticide resistance. Pest Management Science. 2011;67: 886–890.

[11] Weetman D, Djogbenou LS, Lucas E. Copy number variation (CNV) and insecticide resistance in mosquitoes: Evolving knowledge or an evolving problem? Current Opinion in Insect Science. 2018;27: 82–88.

[12] Donnelly MJ, Isaacs AT, Weetman D. Identification, validation, and application of molecular diagnostics for insecticide resistance in malaria vectors. Trends in Parasitology. 2016;32: 197–206.

[13] The *Anopheles gambiae* 1000 Genomes Consortium. Ag1000G phase 2 AR1 data release. MalariaGEN. http://www.malariagen.net/data/ag1000g-phase2-ar1. 2017.

[14] Liu N. Insecticide resistance in mosquitoes: impact, mechanisms, and research directions. Annual Review of Entomology. 2015;60: 537–559.

[15] Weetman D, Mitchell SN, Wilding CS, Birks DP, Yawson AE, Essandoh J, et al. Contemporary evolution of resistance at the major insecticide target site gene Ace-1 by mutation and copy number variation in the malaria mosquito *Anopheles gambiae*. Molecular Ecology. 2015;24: 2656–2672.

[16] Martins AJ, Brito LP, Linss JGB, Rivas GB, Machado R, Bruno RV, et al. Evidence for gene duplication in the voltage-gated sodium channel gene of *Aedes aegypti*. Evolution, Medicine, and Public Health. 2013;2013: 148–160.

[17] Martins WFS, Subramaniam K, Steen K, Mawejje H, Liloglou T, Donnelly MJ, et al. Detection and quantitation of copy number variation in the voltage-gated sodium channel gene of the mosquito *Culex quinquefasciatus*. Scientific Reports. 2017;7: 5821.

[18] Remnant EJ, Good RT, Schmidt JM, Lumb C, Robin C, Daborn PJ, et al. Gene duplication in the major insecticide target site, *Rdl*, in *Drosophila melanogaster*. Proceedings of the National Academy of Sciences. 2013;110: 14705–14710.

[19] Assogba BS, Milesi P, Djogbénou LS, Berthomieu A, Makoundou P, Baba-Moussa LS, et al. The *ace-1* locus is amplified in all resistant *Anopheles gambiae* mosquitoes: fitness consequences of homogeneous and heterogeneous duplications. PLoS Biology. 2016;14: e2000618.

[20] Li X, Schuler MA, Berenbaum MR. Molecular mechanisms of metabolic resistance to synthetic and natural xenobiotics. Annual Review of Entomology. 2007;52: 231–253.

[21] Schmidt JM, Good RT, Appleton B, Sherrard J, Raymant GC, Bogwitz MR, et al. Copy number variation and transposable elements feature in recent, ongoing adaptation at the *Cyp6g1* locus. PLoS Genetics. 2010;6: e1000998.

[22] Itokawa K, Komagata O, Kasai S, Masada M, Tomita T. Cis-acting mutation and duplication: History of molecular evolution in a P450 haplotype responsible for insecticide resistance in *Culex quinquefasciatus*. Insect Biochemistry and Molecular Biology. 2011;41: 503–512.

[23] Raymond M, Berticat C, Weill M, Pasteur N, Chevillon C. Insecticide resistance in the mosquito *Culex pipiens*: what have we learned about adaptation? Genetica. 2001;112: 287–296.

[24] Grigoraki L, Lagnel J, Kioulos I, Kampouraki A, Morou E, Labbé P, et al. Transcriptome profiling and genetic study reveal amplified carboxylesterase genes implicated in temephos resistance, in the Asian Tiger Mosquito *Aedes albopictus*. PLoS Neglected Tropical Diseases. 2015;9: e0003771.

[25] Devonshire AL, Field LM, Foster SP, Moores GD, Williamson MS, Blackman RL. The evolution of insecticide resistance in the peach-potato aphid, *Myzus persicae*. Philosophical Transactions of the Royal Society of London B: Biological Sciences. 1998;353: 1677–1684.

[26] Field LM, Blackman RL, Tyler-Smith C, Devonshire AL. Relationship between amount of esterase and gene copy number in insecticide-resistant *Myzus persicae* (Sulzer). Biochemical Journal. 1999;339: 737–742.

[27] Zimmer CT, Garrood WT, Singh KS, Randall E, Lueke B, Gutbrod O, et al. Neofunctionalization of duplicated P450 genes drives the evolution of insecticide resistance in the brown planthopper. Current Biology. 2018;28: 268 – 274.

[28] Mitchell SN, Rigden DJ, Dowd AJ, Lu F, Wilding CS, Weetman D, et al. Metabolic and target-site mechanisms combine to confer strong DDT resistance in *Anopheles gambiae*. PLoS One. 2014;9: e92662.

[29] Edi CV, Djogbenou L, Jenkins AM, Regna K, Muskavitch MAT, Poupardin R, et al. CYP6 P450 enzymes and ACE-1 duplication produce extreme and multiple insecticide resistance in the malaria mosquito *Anopheles gambiae*. PLoS Genetics. 2014;10: e1004236.

[30] Müller P, Warr E, Stevenson BJ, Pignatelli PM, Morgan JC, Steven A, et al. Field-caught permethrin-resistant Anopheles gambiae overexpress CYP6P3, a P450 that metabolises pyrethroids. PLoS Genetics. 2008;4: e1000286.

[31] Mitchell SN, Stevenson BJ, Müller P, Wilding CS, Egyir-Yawson A, Field SG, et al. Identification and validation of a gene causing cross-resistance between insecticide classes in *Anopheles gambiae* from Ghana. Proceedings of the National Academy of Sciences. 2012;109: 6147–6152.

[32] Stevenson BJ, Bibby J, Pignatelli P, Muangnoicharoen S, O’Neill PM, Lian L, et al. Cytochrome P450 6M2 from the malaria vector *Anopheles gambiae* metabolizes pyrethroids: sequential metabolism of deltamethrin revealed. Insect Biochemistry and Molecular Biology. 2011;41: 492–502.

[33] Chiu T, Wen Z, Rupasinghe SG, Schuler MA. Comparative molecular modeling of *Anopheles gambiae* CYP6Z1, a mosquito P450 capable of metabolizing DDT. Proceedings of the National Academy of Sciences. 2008;105: 8855–8860.

[34] Main BJ, Lee Y, Collier TC, Norris LC, Brisco K, Fofana A, et al. Complex genome evolution in *Anopheles coluzzii* associated with increased insecticide usage in Mali. Molecular Ecology. 2015;24: 5145–5157.

[35] Vontas J, Grigoraki L, Morgan J, Tsakireli D, Fuseini G, Segura L, et al. Rapid selection of a pyrethroid metabolic enzyme CYP9K1 by operational malaria control activities. Proceedings of the National Academy of Sciences. 2018;115: 4619–4624.

[36] Anopheles gambiae 1000 Genomes Consortium. Genetic diversity of the African malaria vector *Anopheles gambiae*. Nature. 2017;552: 96–100.

[37] Schrider DR, Hahn MW, Begun DJ. Parallel evolution of copy-number variation across continents in *Drosophila melanogaster*. Molecular Biology and Evolution. 2016;33: 1308–1316.

[38] Faucon F, Dusfour I, Gaude T, Navratil V, Boyer F, Chandre F, et al. Identifying genomic changes associated with insecticide resistance in the dengue mosquito *Aedes aegypti* by deep targeted sequencing. Genome Research. 2015;25: 1347–1359.

[39] David J, Strode C, Vontas J, Nikou D, Vaughan A, Pignatelli PM, et al. The *Anopheles gambiae* detoxification chip: a highly specific microarray to study metabolic-based insecticide resistance in malaria vectors. Proceedings of the National Academy of Sciences of the United States of America. 2005;102: 4080–4084.

[40] Ding Y, Ortelli F, Rossiter LC, Hemingway J, Ranson H. The *Anopheles gambiae* glutathione transferase supergene family: annotation, phylogeny and expression profiles. BMC genomics. 2003;4: 35.

[41] Riveron JM, Yunta C, Ibrahim SS, Djouaka R, Irving H, Menze BD, et al. A single mutation in the *GSTe2* gene allows tracking of metabolically based insecticide resistance in a major malaria vector. Genome Biology. 2014;15: R27.

[42] Djouaka RF, Bakare AA, Coulibaly ON, Akogbeto MC, Ranson H, Hemingway J, et al. Expression of the cytochrome P450s, CYP6P3 and CYP6M2 are significantly elevated in multiple pyrethroid resistant populations of *Anopheles gambiae* ss. from Southern Benin and Nigeria. BMC genomics. 2008;9: 538.

[43] Kwiatkowska RM, Platt N, Poupardin R, Irving H, Dabire RK, Mitchell S, et al. Dissecting the mechanisms responsible for the multiple insecticide resistance phenotype in *Anopheles gambiae* ss, M form, from Vallée du Kou, Burkina Faso. Gene. 2013;519: 98–106.

[44] Ngufor C, N’Guessan R, Fagbohoun J, Subramaniam K, Odjo A, Fongnikin A, et al. Insecticide resistance profile of *Anopheles gambiae* from a phase II field station in Cové, southern Benin: implications for the evaluation of novel vector control products. Malaria Journal. 2015;14: 464.

[45] Fossog Tene B, Poupardin R, Costantini C, Awono-Ambene P, Wondji CS, Ranson H, et al. Resistance to DDT in an urban setting: common mechanisms implicated in both M and S forms of *Anopheles gambiae* in the city of Yaoundé Cameroon. PloS One. 2013;8: e61408.

[46] Müller P, Donnelly MJ, Ranson H. Transcription profiling of a recently colonised pyrethroid resistant *Anopheles gambiae* strain from Ghana. BMC genomics. 2007;8: 36.

[47] Nikou D, Ranson H, Hemingway J. An adult-specific CYP6 P450 gene is overexpressed in a pyrethroid-resistant strain of the malaria vector, *Anopheles gambiae*. Gene. 2003;318: 91–102.

[48] Thomsen EK, Strode C, Hemmings K, Hughes AJ, Chanda E, Musapa M, et al. Underpinning sustainable vector control through informed insecticide resistance management. PLoS One. 2014;9: e99822.

[49] Lumjuan N, Rajatileka S, Changsom D, Wicheer J, Leelapat P, Prapanthadara L, et al. The role of the *Aedes aegypti* Epsilon glutathione transferases in conferring resistance to DDT and pyrethroid insecticides. Insect Biochemistry and Molecular Biology. 2011;41: 203–209.

[50] Ibrahim SS, Amvongo-Adjia N, Wondji MJ, Irving H, Riveron JM, Wondji CS. Pyrethroid resistance in the major malaria vector *Anopheles funestus* is exacerbated by overexpression and overactivity of the P450 *CYP6AA1* across Africa. Genes. 2018;9: 140.

[51] Riveron JM, Ibrahim SS, Chanda E, Mzilahowa T, Cuamba N, Irving H, et al. The highly polymorphic CYP6M7 cytochrome P450 gene partners with the directionally selected CYP6P9a and CYP6P9b genes to expand the pyrethroid resistance front in the malaria vector *Anopheles funestus* in Africa. BMC Genomics. 2014;15: 817.

[52] Rodpradit P, Boonsuepsakul S, Chareonviriyaphap T, Bangs MJ, Rongnoparut P. Cytochrome P450 genes: molecular cloning and overexpression in a pyrethroid-resistant strain of *Anopheles minimus* mosquito. Journal of the American Mosquito Control Association. 2005;21: 71–79.

[53] Duangkaew P, Pethuan S, Kaewpa D, Boonsuepsakul S, Sarapusit S, Rongnoparut P. Characterization of mosquito CYP6P7 and CYP6AA3: differences in substrate preference and kinetic properties. Archives of Insect Biochemistry and Physiology. 2011;76: 236–248.

[54] Abyzov A, Urban AE, Snyder M, Gerstein M. CNVnator: an approach to discover, genotype, and characterize typical and atypical CNVs from family and population genome sequencing. Genome Research. 2011;21: 974–984.

[55] Miles A, Iqbal Z, Vauterin P, Pearson R, Campino S, Theron M, et al. Indels, structural variation, and recombination drive genomic diversity in *Plasmodium falciparum*. Genome rResearch. 2016;26: 1288–1299.

[56] Alexa A, Rahnenfuhrer J. topGO: Enrichment analysis for Gene Ontology. R package version 2.14.0. 2010.

[57] Klaus B, Strimmer K. fdrtool: Estimation of (local) false discovery rates and higher Criticism. R package version 1.2.15. 2015.

[58] Sabeti PC, Reich DE, Higgins JM, Levine HZP, Richter DJ, Schaffner SF, et al. Detecting recent positive selection in the human genome from haplotype structure. Nature. 2002;419: 832– 837.

[59] R Development Core Team. R: A Language and Environment for Statistical Computing. 2008.

[60] Menelaou A, Marchini J. Genotype calling and phasing using next-generation sequencing reads and a haplotype scaffold. Bioinformatics. 2013;29: 84–91.

